## Supplementary Figures for "Ubiquitination-mediated Golgi-to-endosome sorting determines the toxin-antidote duality of *wtf* meiotic drivers"

Supplementary Table 1–3 are provided as separate Excel files.

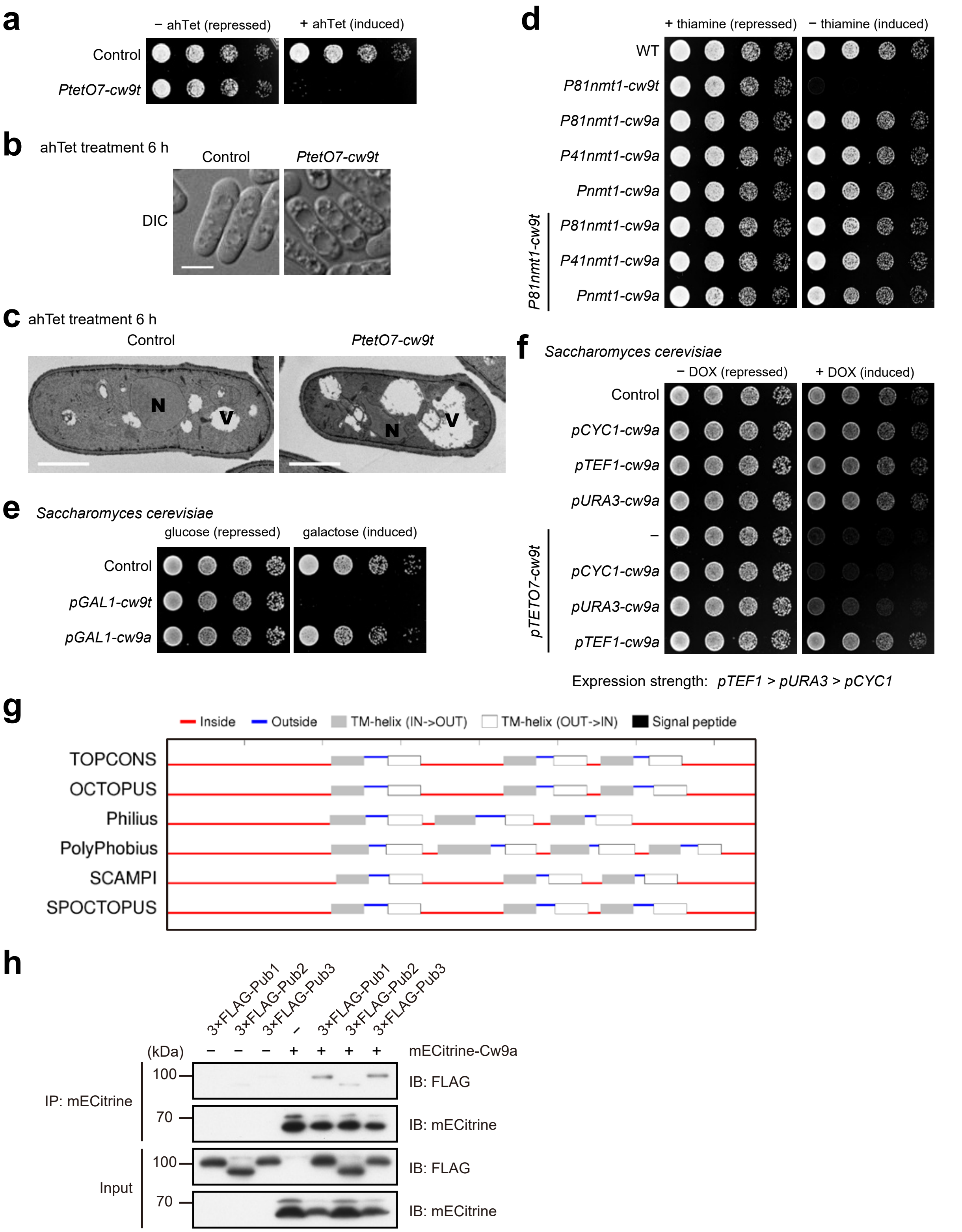

**Supplementary Figure 1. The toxicity of Cw9t and the detoxification activity of Cw9a.**

**a** Cw9t expressed from the *PtetO7* promoter caused toxicity in vegetative *S. pombe* cells.

**b** Cw9t expressed from the *PtetO7* promoter caused vacuole enlargement in vegetative *S. pombe* cells. DIC, differential interference contrast. Bar, 5 μm.
**c** Electron microscopy images of control cells containing an empty vector and cells expressing Cw9t from the *PtetO7* promoter. N, nucleus; V, vacuole. Bar, 2 μm.
**d** Cw9a neutralized the toxicity of Cw9t expressed from the *P81nmt1* promoter.
**e** Cw9t was toxic to vegetative *S. cerevisiae* cells. *pGAL1* is a galactose-inducible promoter.
**f** Cw9a neutralized the toxicity of Cw9t in a dose-dependent manner in *S. cerevisiae*. *pTETO7* promoter is a doxycycline (DOX)-inducible promoter and *pTEF1*, *pURA3*, and *pCYC1* are three constitutive promoters with different strengths.
**g** Transmembrane topology of Cw9a was predicted using the TOPCONS web server (https://topcons.cbr.su.se/pred/). TOPCONS is a consensus-based method integrating the results of five different topology prediction algorithms. The topology predictions using the individual methods and the consensus prediction of TOPCONS 2.0 are shown. Inside indicates the cytosolic side of the membrane and outside indicates the non-cytosoilic side of the membrane.
**h** Pub1, Pub2, and Pub3 were coimmunoprecipitated with Cw9a. All proteins were expressed from the *P41nmt1* promoter.

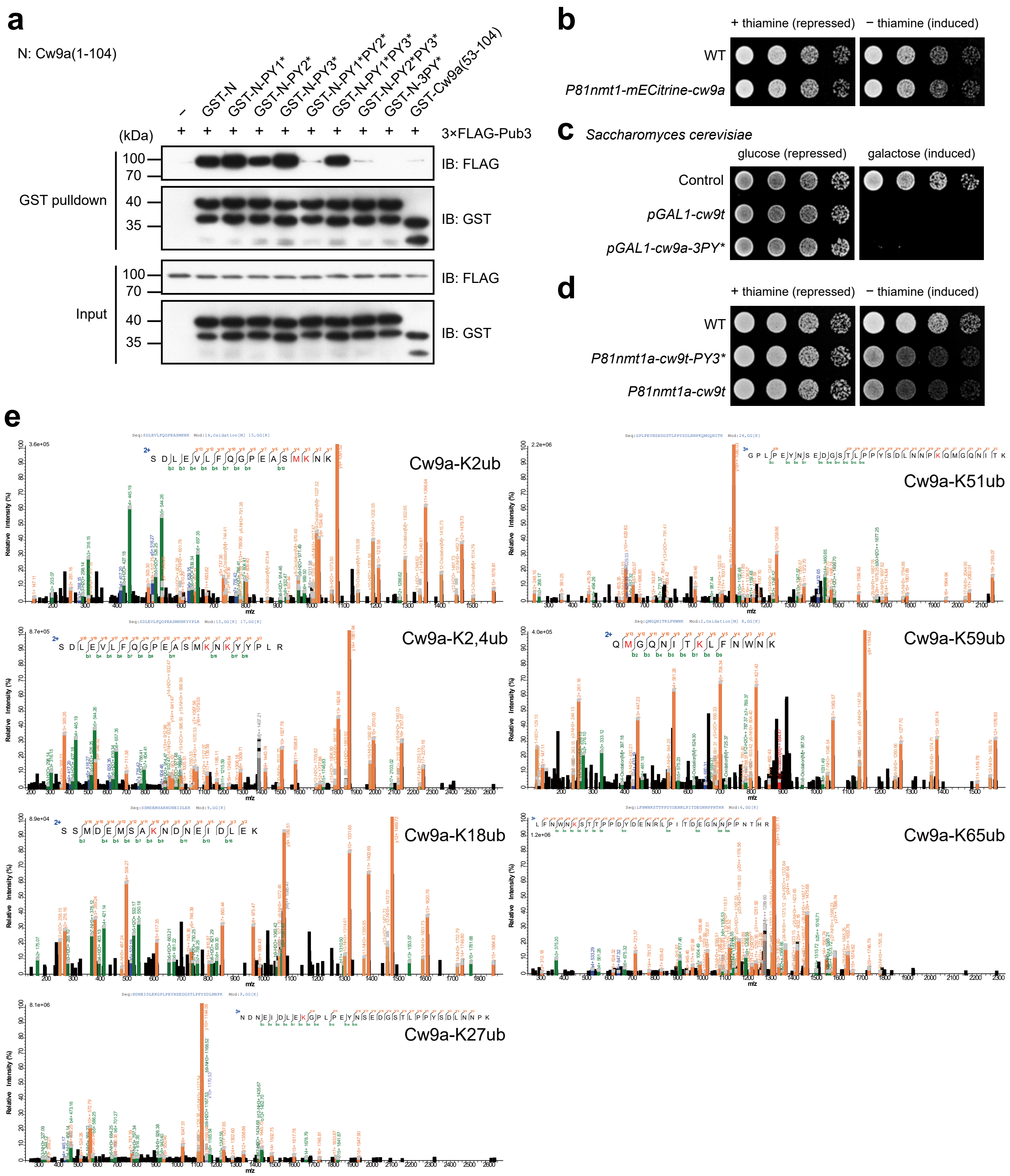

**Supplementary Figure 2. PY motif-mediated ubiquitin ligase binding drives Cw9a ubiquitination.**

**a** GST pulldown assay showed that PY motifs in the N-terminal cytosolic tail of Cw9a mediate Pub3 binding. Lysates of *E. coli* cells expressing GST-tagged Cw9a N-terminal fragments were mixed with the lysate of *S. pombe* cells expressing 3xFLAG-Pub3 and pulldown was performed using glutathione beads.

**b** mECitrine-Cw9a showed no toxicity to vegetative *S. pombe* cells.

**c** Cw9a-3PY* was toxic to vegetative *S. cerevisiae* cells when expressed from the *pGAL1* promoter.
**d** Mutating PY3 in Cw9t did not affect its toxicity. Expression was under the control of the *P81nmt1a* promoter, which is an attenuated version of the *P81nmt1* promoter.
**e** Representative mass spectrometry spectra of ubiquitinated peptides from in vitro ubiquitinated GST-Cw9a(1-104).

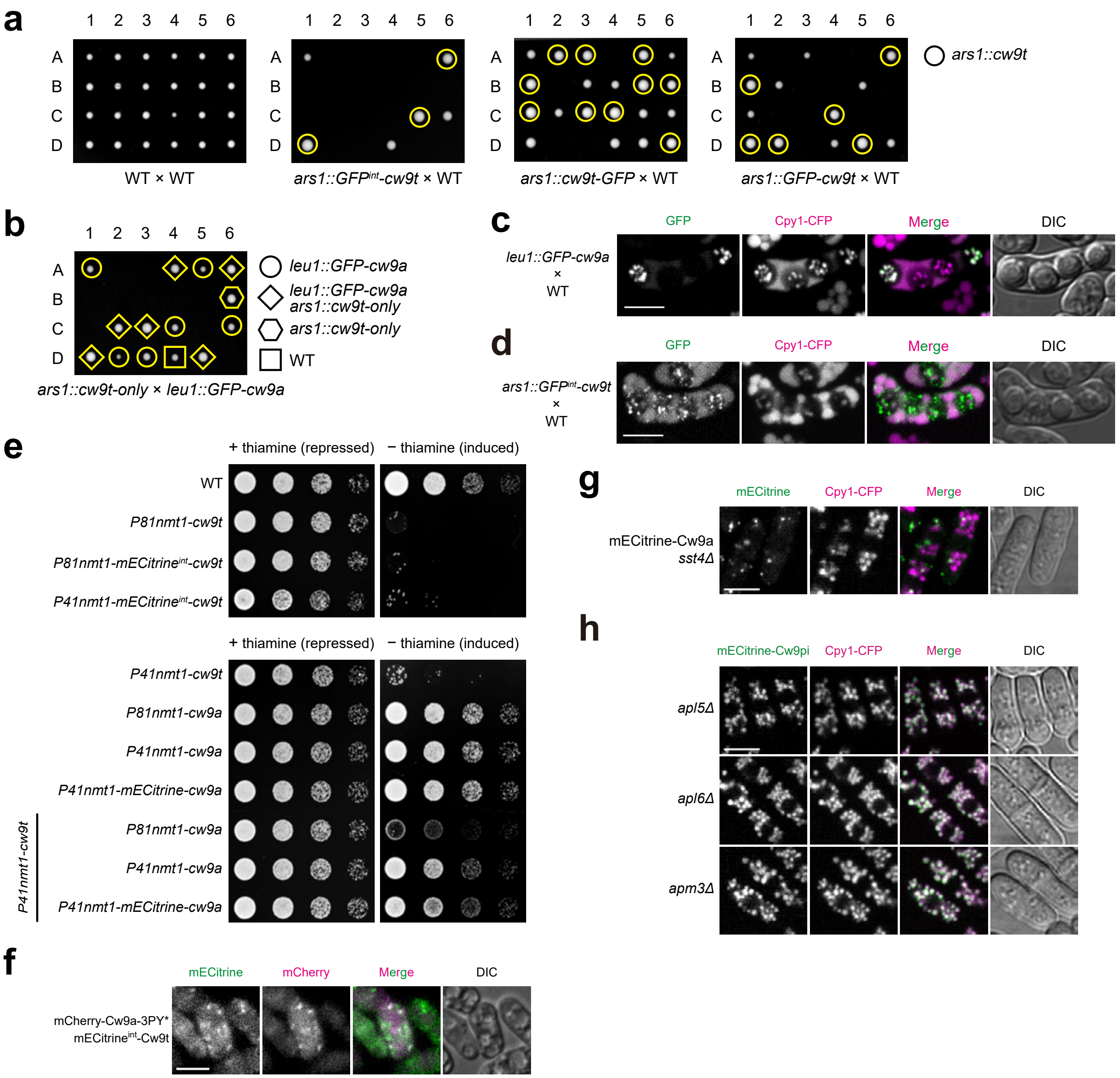

**Supplementary Figure 3. Subcellular localizations of Cw9a and Cw9t.**

**a** When under the control of the native promoter, internally GFP-tagged Cw9t exhibited a stronger spore killing than C-terminally and N-terminally tagged Cw9t. A *ura4*$^{+}$ marker linked to the Cw9t coding sequence caused faster growth of the surviving colonies containing the Cw9t coding sequence.

**b** GFP-tagged Cw9a under the control of the native promoter was able to neutralize the spore killing activity of Cw9t.

**c** Cw9a localized to the vacuole in spores. Cpy1-CFP is a vacuole lumen marker. Bar, 5 μm.

**d** Cw9t localized to cytoplasmic puncta outside of vacuoles in spores. Bar, 5 μm.

**e** In vegetative cells, mECitrine-tagged Cw9t caused growth inhibition and mECitrine-tagged Cw9a conferred protection against Cw9t.

**f** Cw9a-3PY* co-localized with Cw9t. Cw9a-3PY* was N-terminally tagged and Cw9t was internally tagged. Bar, 5 μm.

**g** The vacuolar targeting of Cw9a was abolished in *sst4Δ* cells. Cw9a was N-terminally tagged. Bar, 5 μm.
**h** The vacuole lumen localization of Cw9a was unaffected in AP3 deficient cells. Cw9a was N-terminally tagged. Bar, 5 μm.

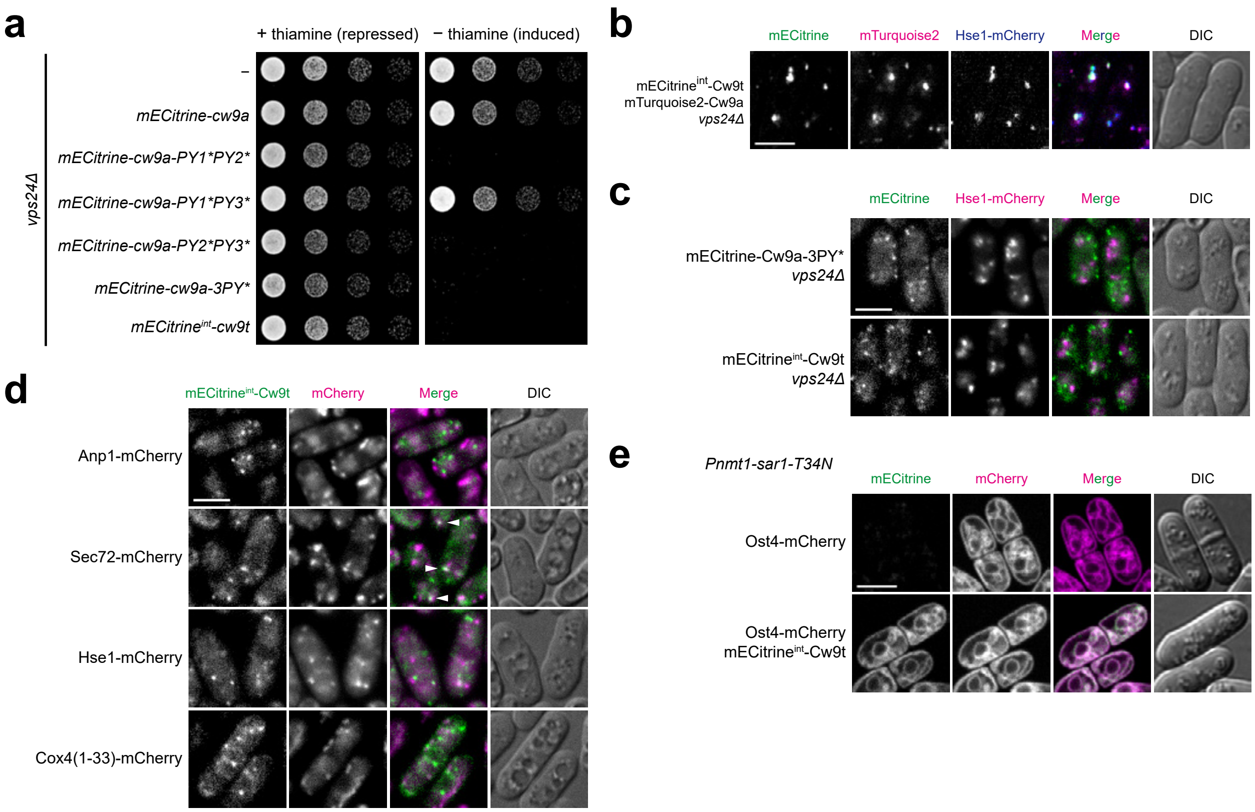

**Supplementary Figure 4. ESCRT-mediated trafficking of Cw9a and the trafficking route of Cw9t.**

**a** Cw9t and the *PY1*PY2**, *PY2*PY3**, and *3PY** mutants of Cw9a exhibited toxicity in the *vps24Δ* background. Expression was under the control of the *P81nmt1* promoter.

**b** Cw9t localized to the endosome when co-expressed with Cw9a in the *vps24Δ* background. Bar, 5 μm.

**c** Cw9a-3PY* and Cw9t did not co-localize with the endosome marker Hse1 in *vps24Δ* cells. Bar, 5 μm.

**d** Co-localization analysis of Cw9t with markers of different subcellular organelles. Internally tagged Cw9t was expressed from the *P81nmt1* promoter. Anp1-mCherry is a cis-Golgi marker. Sec72-mCherry is a TGN marker. Hse1-mCherry is an endosome marker. Cox4(1-33)-mCherry is a mitochondria marker. Arrowheads indicate overlapping signals of Cw9t and Sec72. Bar, 5 μm.

**e** Overexpression of Sar1-T34N changed the localization of Cw9t to the ER. Sar1-T34N was expressed from the *Pnmt1* promoter. Internally tagged Cw9t was expressed from the *P81nmt1* promoter. Ost4-mCherry is an ER marker. Bar, 5 μm.

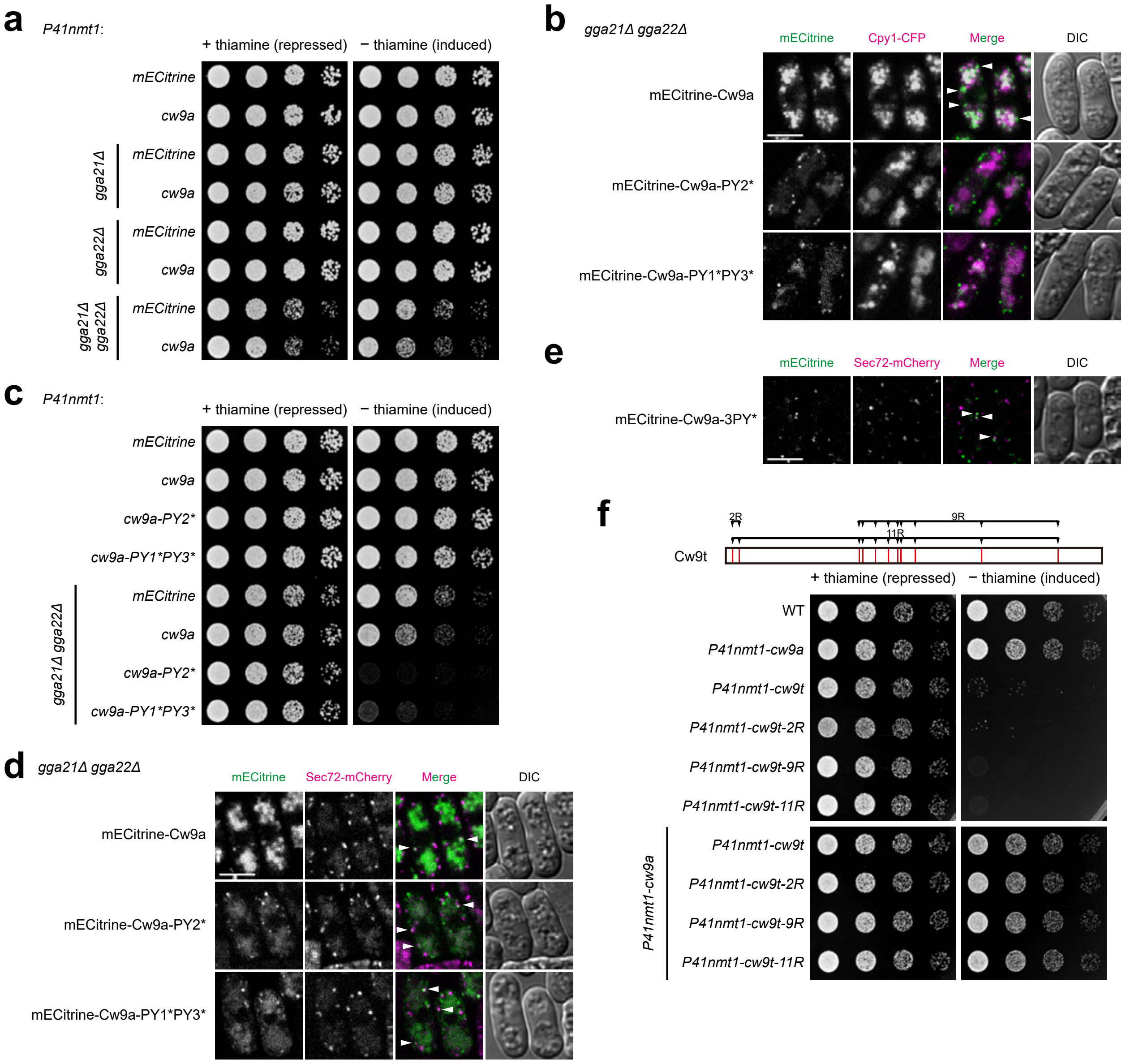

**Supplementary Figure 5. GGA proteins promote the TGN-to-endosome trafficking of Cw9a.**

**a** Cw9a caused a mild growth inhibition in the *gga21Δ gga22Δ* background.

**b** Localization of Cw9a and its *PY2** and *PY1*PY3** mutants in the *gga21Δ gga22Δ* background. Expression was under the control of the *P41nmt1* promoter. Arrowheads indicate Cw9a signals outside of vacuoles. Bar, 5 μm.

**c** The *PY2** and *PY1*PY3** mutants of Cw9a exhibited strong toxicity in the *gga21Δ gga22Δ* background.

**d** Cytoplasmic puncta of Cw9a and its *PY2** and *PY1*PY3** mutants partially co-localized with the TGN marker Sec72 in the *gga21Δ gga22Δ* background. Expression was under the control of the *P41nmt1* promoter. Arrowheads indicate signals overlapping with Sec72-mCherry. Bar, 5 μm.

**e** Cw9a-3PY* partially co-localized with the TGN marker Sec72. N-terminally tagged Cw9a-3PY* was expressed from the *P81nmt1* promoter. Arrowheads indicate signals overlapping with Sec72-mCherry. Bar, 5 µm.
**f** Mutating all cytosol-facing lysines of Cw9t to arginines did not affect its toxicity or the neutralization of its toxicity by Cw9a. Expression was under the control of the *P41nmt1* promoter.

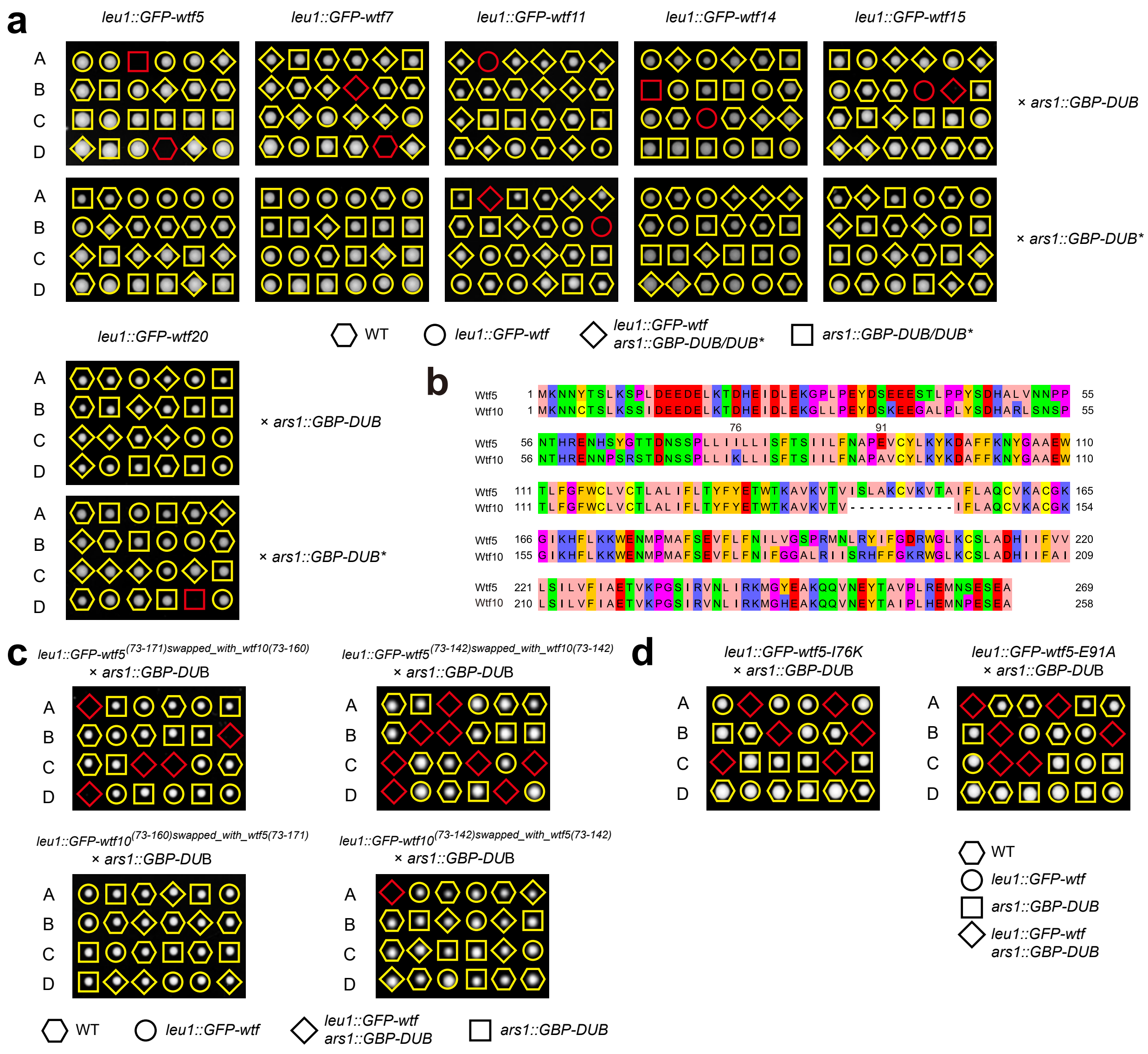

**Supplementary Figure 6. The antidote products of certain *S. pombe wtf* genes did not exhibit toxicity when tethered to a DUB.**

**a** The antidote products of six *S. pombe wtf* genes of the *S. pombe* reference genome were not rendered toxic by artificial DUB tethering. Experiments were performed as in Fig. 6a.

**b** The sequence alignment of Wtf5 and Wtf10*.* The positions of the two residues (Ile76 and Glu91) affecting toxicity are shown.

**c** Sequence swapping between Wtf5 and Wtf10. Experiments were performed as in Fig. 6a.

**d** Either an I76K mutation or an E91A mutation rendered Wtf5 toxic in the DUB tethering analysis. Experiments were performed as in Fig. 6a.

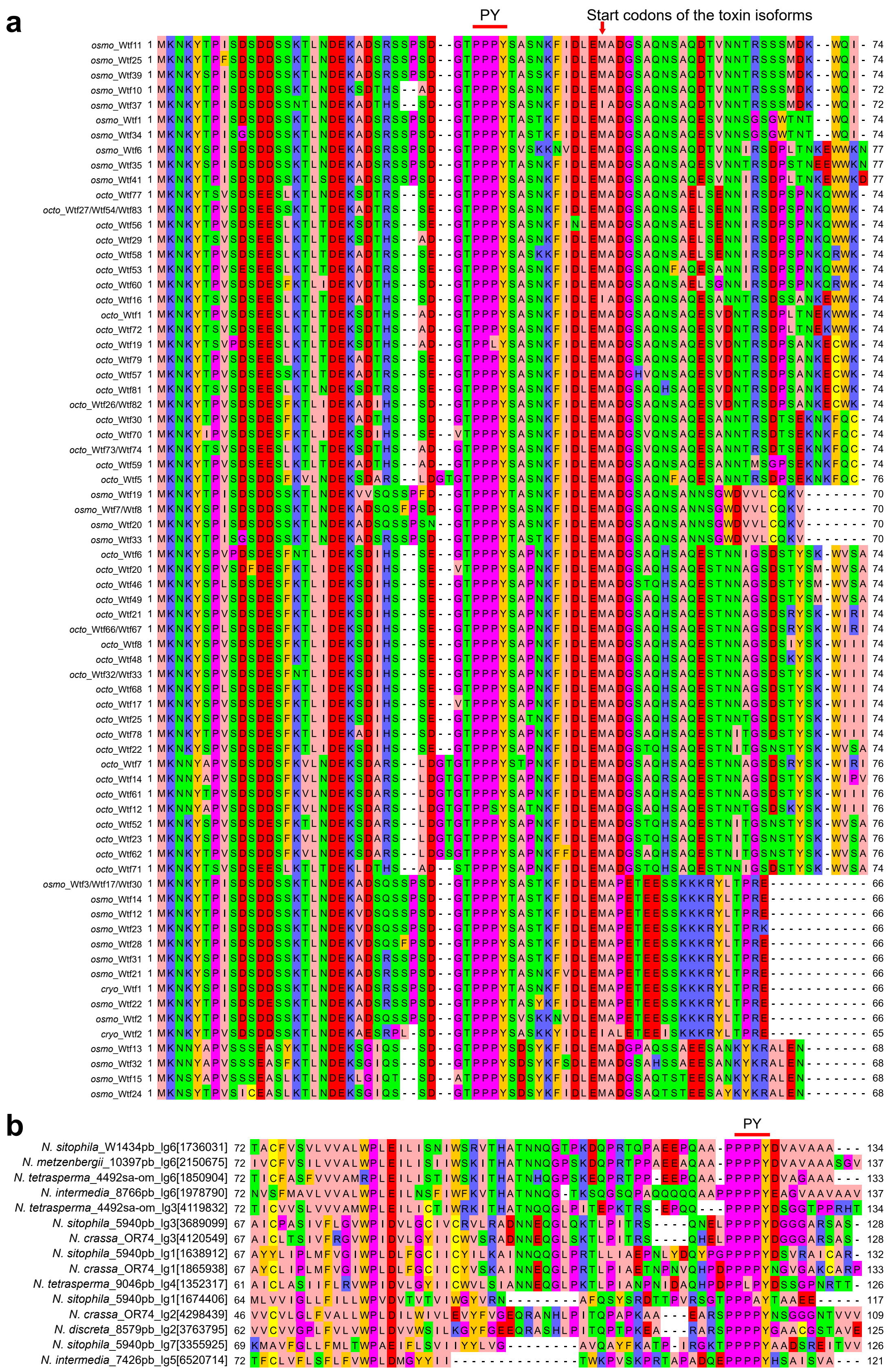

**Supplementary Figure 7. PY motifs are present in the protein products of non-*S.***

***pombe* KMDs.**

**a** Multiple sequence alignment of the N-terminal cytosolic tails of the antidote products encoded by *wtf* genes in *S. octosporus*, *S. osmophilus*, and *S. cryophilus*. Proteins sharing the same sequence in the N-terminal cytosolic tails are shown in a single row. The conserved PY motif is highlighted.

**b** Multiple sequence alignment of the C-terminal region of proteins encoded by representative *Spk-1* family genes. The conserved PY motif is highlighted.
